# Systematic Evaluation of Lecithin:Cholesterol Acyltransferase Binding Sites in Apolipoproteins via Peptide Based Nanodiscs: Regulatory Role of Charged Residues at Positions 4 and 7

**DOI:** 10.1101/2023.12.20.572495

**Authors:** Akseli Niemelä, Artturi Koivuniemi

## Abstract

Lecithin:cholesterol acyltransferase (LCAT) exhibits α-activity on high-density and β-activity on low-density lipoproteins. However, the molecular determinants governing LCAT activation by different apolipoproteins remain elusive. Uncovering these determinants would offer the opportunity to design and explore advanced therapies against dyslipidemias. Here, we have conducted coarse-grained and all-atom molecular dynamics simulations of LCAT with nanodiscs made with α-helical amphiphilic peptides either derived from apolipoproteins A1 and E (apoA1 and apoE) or apoA1 mimetic peptide 22A that was optimized to activate LCAT. This study aims to explore what drives the binding of peptides to our previously identified interaction site in LCAT. We hypothesized that this approach could be used to screen for binding sites of LCAT in different apolipoproteins and would provide insights to differently localized LCAT activities. Our screening approach was able to discriminate apoA1 helixes 4, 6, and 7 as key contributors to the interaction with LCAT supporting the previous research data. The simulations provided detailed molecular determinants driving the interaction with LCAT: the formation of hydrogen bonds or salt bridges between peptides E4 or D4 and LCAT S236 or K238 residues. Additionally, salt bridging between R7 and D73 was observed, depending on the availability of R7. Expanding our investigation to diverse plasma proteins, we detected novel LCAT binding helixes in apoL1, apoB100, and serum amyloid A. Our findings suggest that the same binding determinants, involving E4 or D4 -S236 and R7-D73 interactions, influence LCAT β-activity on low-density lipoproteins, where apoE and or apoB100 are hypothesized to interact with LCAT.

## INTRODUCTION

Reverse cholesterol transport (RCT) is driven by high-density lipoprotein (HDL) particles in plasma. The smallest type of HDL particle is pre-β1 HDL also known as nascent HDL (nHDL), which consists of two apolipoproteins surrounding a lipid bilayer in a double belt configuration [1,2]. It is formed when lipid poor apolipoprotein A1 (apoA1) attains a lipid bilayer of phospholipids and cholesterol from ATP-binding cassette transporters (ABCA1) of peripheral cells. As lecithin:cholesterol acyltransferase (LCAT) encounters nHDL in plasma, it binds to it and separates an acyl chain from a phospholipid resulting in a lysolipid [3] and LCAT-bound acyl chain. LCAT then attaches the acyl chain to cholesterol resulting in a cholesteryl ester (CE). The CEs travel to the center of the bilayer forming a hydrophobic core, which turns the discoidal nHDL to a spherical mature HDL particle, also known as α-HDL. The lipid composition of mature HDL is further modified by LCAT and cholesteryl ester transfer protein (CETP), which mainly trades the triglycerides of very low-density lipoprotein particles for the CEs of HDL. Finally, upon reaching the liver, the cholesterol and CEs of HDL get selectively deposited by the scavenger receptor B1 to hepatocytes for further metabolism and excretion, while the apoA1 gets released as lipid poor apoA1, completing the traditional model of the RCT cycle.

As RCT drives cholesterol from peripheral cells to the liver, it has been hypothesized to prevent the formation of atherosclerosis, where CEs accumulate to the walls of arterial vessels. Indeed, an increase in HDL cholesterol (HDL-C) is negatively correlated with cardiovascular disease incidence [4]. This led to the development of HDL-C raising therapies whose goal was to prevent atherosclerotic cardiovascular disease. The most successful of these was CETP-inhibitor anacetrapib, whose mechanism of action is preventing CE escape from HDL to LDL leading to improved HDL-C levels. However, its commercialization was ultimately ceased and its effectiveness in clinical trials was instead attributed to the decrease of non-HDL-C levels [5]. Furthermore, a newer model of RCT posits that the majority of cholesterol transport occurs with free cholesterol through passive reversible pathways instead of CEs in transporters [6]. This reversibility means that when HDL-C is increased it may also lead to increased cholesterol influx to peripheral cells, negating its atheroprotective effect.

Although the new RCT model diminishes the role of LCAT, LCAT is still vital for the formation of healthy lipoprotein particles and thus RCT as a whole. This can be seen in patients whose LCAT activity is inhibited by mutations. Although LCAT primarily acts on HDL via apoA1, called α-activity, it also has some activity on LDL particles, called β-activity [7]. Apolipoprotein E (apoE) has been identified as the primary cofactor in β-activity [8]. LCAT mutations have been classified to two groups based on whether α-activity or both α- and β-activities is lost [9]. A loss of α-activity leads to cholesterol accumulating in cornea, causing a condition called fish-eye disease (FED). FED has also been linked to atherosclerosis [10]. A loss of both α- and β-activities causes familial LCAT deficiency (FLD) and it leads to more severe complications like renal failure. Interestingly however, it is seemingly atheroprotective. This property of FLD is suggested to be caused by the rapid metabolism of dysfunctional CE poor LDL, which then prevents CE accumulation to arterial walls [11,12].

Based on these findings, we are interested in exploring if the β-activity of LCAT can be selectively inhibited without harming its α-activity, and whether this would be beneficial against atherosclerosis. We hypothesize that β-activity may have diminished the efficacy and safety of recombinant human LCAT (MEDI6012) in its recent phase 2b clinical trial [13]. A challenge for the first question is the fact that no such LCAT mutations have been discovered whereas several FED and FLD causing mutations have been. Either the selective loss of β-activity leads to neutral or benign outcomes and thus it hasn’t been noticed, or the mechanism for α- and β-activities has a common mechanism to which α-activity imposes additional conditions. Thus, any mutation harming β-activity would also harm α-activity, but not vice versa. A challenge for the second question is that a loss of β-activity may lead to the formation of nephrotoxic lipoprotein X (LpX), as increased LpX is associated with FLD but not with FED [14]. However, as LCAT and apoA1 are able to remodel LpX to HDL-like particles, any LCAT activity may protect against it. Given that β-inhibition would still leave the CETP pathway for CEs to enter LDL intact, anacetrapib could provide a synergistic effect. Regardless, passive free cholesterol transport pathways from cholesterol rich HDL remains, which in the context of RCT may only be solved by increasing α-activity as well.

We previously investigated the interaction between LCAT and nHDL-like nanodiscs formed with apoA1 mimetic peptides through a combination of computational and experimental methods [15]. These peptides, specifically 22A and some of its mutants, are based on the repeating amphiphilic helixes of apoA1. Coarse-grained (CG) molecular dynamics (MD) simulations indicated that LCAT in its open active form takes a spatially restricted position in the rim of the nanodiscs. At this position, which matched electron microscopy (EM) images, the mimetic peptides bound to the cavity between LCAT’s lid and membrane binding domain (MBD) with their hydrophilic side. After measuring LCAT’s acyltransferase activity experimentally with the different peptide variants, we noted that their in vitro activities were in line with their in silico occupancies in the binding site, suggesting that the in silico approach can be utilized to screen for potential LCAT activating peptides and binding sites in apolipoproteins.

Our aim was to expand on our previous CG MD simulations to find LCAT’s binding site on apoA1 and apoE, which could then be exploited in the design of mutants that selectively inhibit the β-activity of LCAT. Although other apolipoproteins may end up being better targets for inhibition than apoE, it is a suitable starting point to characterize the common residues or residue types with which apolipoproteins interact with LCAT. We were successful in finding binding sites on apoA1 which agreed with existing experimental data. Furthermore, we identified an amino acid pattern helical peptides utilize in binding to LCAT. The pattern was further utilized in screening for LCAT binding helixes in other apolipoproteins and proteins associated with lipoproteins. These findings pave the way for discovering an LCAT therapy with a novel mechanism of action and for elucidating the LCAT-apolipoprotein interaction.

## METHODS

### System preparation and running parameters

Coarse-grained Martini 3 nanodisc systems with LCAT were all prepared similarly for the GROMACS simulation package [16,17]. This refers to the systems labelled A1-8, E1-8, R1-5, M1-2 and N1-27. The preparation of systems L1-2 is described in our previous publication, which after equilibration were equal to the new systems [15]. The same model for coarse-grained open LCAT was used as previously, which was based on the X-ray crystallographic structure of PDB ID 6MVD [18]. All peptides were first built in atomistic resolution with the protein builder of Molefacture in VMD preset to an α-helical secondary structure [19]. The peptides were coarse-grained with Martinize2 [20], but unlike LCAT whose secondary structure was determined with DSSP [21,22] and restrained with an elastic network, the secondary structure of the first and last residues of the peptides was set to random coils, while the rest of the structure was set to α-helical (except N3 where residue 12 was set to random coil). The CG models for DMPC and water were provided in the Martini 3 release.

With the attained coordinates and topologies, the protocol to prepare these systems was as follows. With gmx insert-molecules, where the preceding gmx refers to a GROMACS function included in the package, 28 peptides were placed randomly to a box of 13 x 13 x 13 nm^3^ followed by 200 randomly placed DMPC. The peptide-lipid system was energy minimized and ran with a semi-isotropic Parrinello-Rahman barostat [23] targeting 1 bar in both directions but with a 1*10^−4^ bar^−1^ and 3*10^−4^ bar^−1^ compressibility for x/y and z directions respectively for 100 ps with a 20 fs time step. The pressure coupling time constant was 12 ps. The temperature was coupled with a v-rescale thermostat for 320 K with a 1 ps time constant [24]. Neighbour searching was performed with Verlet every 20 steps and periodic boundary conditions were applied to all sides. Van der Waals interactions were handled with a 1 nm cut-off and with a potential shift Verlet modifier, and electrostatic interactions with the reaction field method using a 1.1 nm Coulomb cut-off and a relative dielectric constant of 15 and an infinite relative dielectric constant of the reaction field. This system forces the beads together to form a flat surface. The flat surface is placed to the centre of a 20 x 20 x 20 nm^3^ box and open LCAT placed on one side of the surface a few nanometers away from the surface. The system is solvated with ∼64 000 water beads via gmx solvate and energy minimized. Finally, the system is set to run for 20 µs using the same properties as above except with an isotropic barostat set to 1 bar with a 3*10^−4^ bar^−1^ compressibility. As some peptides were highly hydrophilic, they were not able to bind to the DMPC bilayer at all, and thus their runs were cancelled. For the N1-24 systems binding to LCAT was analysed at 8 µs and the run discontinued if the binding was poor.

To test how well LCAT would bind to full length apoA1 if its helix 4 was bound to the same binding site, a CG system was built using the ideal atomistic resolution model of apoA1 provided in e.g. PDB ID 2MSC. The system is labelled S1. The model was truncated to only include residues 44-241, and the N- and C-termina were covalently attached together to form a cyclic protein. With Martinize2 the secondary structure was defined manually to include a random coil kink at each proline residue and the double glycine residue of helix 7. Separately, 120 DMPC were forced together into a dense cube as described above but with an isotropic barostat. The circular cyclic apoA1 was placed to the centre of a 20 x 20 x 20 nm^3^ box and the DMPC cube inside it. The backbone beads of apoA1 were position restrained and the system was run with the same conditions as above for 90 ns to ensure the lipid bilayer flattened inside the circle. LCAT was placed to have helix 4 at the binding site and a 10 µs run was conducted without position restraints.

To investigate how 22A and 22A-R7Q behave in the binding site in all-atom (AA) resolution, first, a system using the same 1:7 peptide:lipid ratio was prepared in CG with 9 peptides and 60 DMPC and open LCAT with the same technique as above. The systems were equilibrated for 1 µs and run until three frames at least 0.5 µs apart with a peptide bound to LCAT were found. Coordinates were mapped to atomistic resolution using CHARMM-GUI [25,26]. For other peptides (I3-7 and B1) the same strategy was used to find one frame. Topologies based on the CHARMM36 force field were obtained [27], and in order to increase sampling hydrogen mass repartitioning (HMR) was used [28]. Atomistic open LCAT structure based on PDB ID 6MVD prepared with the same protocol as in our previous publication was superimposed on the backmapped LCAT, whose natural secondary structure details had been lost in the conversion [29,30]. The peptide-lipid-LCAT complex was placed to the centre of a 12 x 12 x 12 nm^3^ box. The system was solvated with ∼48000 water molecules based on the TIP3P model [31]. Ions were added to neutralize the systems. The systems were first equilibrated using 10 kJ/mol nm^2^ position restraints on the peptide α carbons and LCAT backbone atoms for 1 ns with the Berendsen barostat. The production systems without restraints were run for 300 ns with a 4 fs time step, afforded by HMR. One 22A system was run to 600 ns. The temperature was handled with a v-rescale thermostat set to 310 K with a 1 ps time constant and the pressure with a Parinello-Rahman barostat set to 1.0 bar and a 4.5 * 10^−5^ bar^−1^ compressibility and a 5 ps time constant. A cut-off scheme was used for Van der Waals interactions with a 1.2 nm cut-off and a force switch modifier at 1.0 nm, and Particle Mesh Ewald was used for electrostatics with a 1.2 nm cut-off. Hydrogen bonds were constrained with LINCS. For evaluating solvent accessible surface area differences, an atomistic closed LCAT system without a nanodisc was run with the same settings but in a 8.4 x 8.4 x 8.4 nm^3^ box for 300 ns. A closed LCAT model based on the PDB ID 5TXF structure was prepared with the same protocol and run with the same topology as open LCAT [32].

### Analysis

For the CG systems, peptide binding to a specific site on LCAT was measured with a custom script. At each frame the nearest peptide to the site was identified by having the smallest combined distance between backbone beads of peptide residue 2 to LCAT residue G226, peptide residue 11 to the middle point between LCAT residues W48 and I237, and peptide residue 20 to LCAT W48. If the distances are less than 1 nm, 1 nm and 2 nm respectively for the nearest peptide, LCAT is considered to be occupied that frame. Based on this condition we obtain occupancy, which is the percentage of occupied frames to the total number of analysed frames. The analysis was set to start at 1 µs or when LCAT found its position on the rim of the nanodisc. To attain standard deviations the data was divided to three equal sized blocks (i.e. data over 18 µs was divided to blocks of 6 µs) and the standard deviation between the mean values of each block was measured. For L1-2 each simulation was treated as a block.

As the CG systems also showcase peptide behaviour on nanodiscs generally, we measured their dimerization tendencies with 2D density plots of peptide-peptide residue 13 distances at 0-2 nm vs peptide-peptide residue 1 to 21 vector angles at 0-180 degrees. This was measured for all peptide pair combinations at each frame. As hydrophilic peptides had a tendency to escape the bilayer to the water phase, the percentage of dissociated peptides was counted by measuring the minimum distance between a peptide’s residue 13 backbone bead and all the C3A tail beads of the DMPCs. If the minimum distance was over 2 nm the peptide was considered dissociated. Standard deviation was measured by dividing the data to three equal blocks after equilibration. These analyses were set to begin after 1 µs, and standard deviations were measured by dividing the data to three equal blocks.

The mean hydrophilicity of the peptides was estimated with the hydrophilicity values of Hopp and Woods [33]. Additionally, mean helical penalties were calculated by using the values of Pace and Scholtz [34]. A value of 0.75 and 2 kJ/mol were determined to be good cut-offs for hydrophilicity and helical penalty respectively when screening for helical peptides with sufficient binding to the lipid bilayer. PEP-FOLD3 was used to determine their secondary structures [35]. Hydrophobic moments for 22 amino acid long sequences were calculated with the HELIQUEST server but weren’t used in screening [36].

Atomistic simulations were inspected visually and by counting the number of hydrogen bonds between the relevant residues of the peptide and LCAT with gmx hbond using standard settings after a stable conformation was reached. Per residue solvent accessible surface area differences between open and closed LCAT was calculated by subtracting the gmx sasa results, attained with standard settings, of the 600 ns LCAT and 22A based nanodisc simulation from the closed LCAT in water simulation.

## RESULTS AND DISCUSSION

All simulated systems are shown in Table 1. The sequences for the peptides are collected in Table S1.

**Table 1.** All simulated systems and their identifiers.

| ID | res | peptide | composition | runtime |
| --- | --- | --- | --- | --- |
| A1 | CG | apoA1 helix 1 | 28 peptide + 200 DMPC + 1 open LCAT in 20x20x20 nm<sup>3</sup> | 20 us |
| A2 | CG | apoA1 helix 2 | " | 20 us |
| A3 | CG | apoA1 helix 4 | " | 20 us |
| A4 | CG | apoA1 helix 5 | " | 20 us |
| A5 | CG | apoA1 helix 6 | " | 20 us |
| A6 | CG | apoA1 helix 7 | " | 20 us |
| A7 | CG | apoA1 helix 8 | " | 20 us |
| A8 | CG | apoA1 helix 10 | " | 20 us |
| E1 | CG | apoE helix 1 | " | 20 us |
| E2 | CG | apoE helix 2 | " | cancelled |
| E3 | CG | apoE helix 3 | " | 20 us |
| E4 | CG | apoE helix 4 | " | 20 us |
| E5 | CG | apoE helix 5 | " | cancelled |
| E6 | CG | apoE helix 6 | " | 20 us |
| E7 | CG | apoE helix 7 | " | cancelled |
| E8 | CG | apoE helix 8 | " | cancelled |
| S1 | CG | cyclic truncated apoA1 | 1 peptide + 120 DMPC + 1 open LCAT in 20x20x20 nm<sup>3</sup> | 10 us |
| S2 | CG | apoA1 helix 4 and apoA1 helix 6 | 14+14 peptide + 200 DMPC + 1 open LCAT in 20x20x20 nm<sup>3</sup> | 10 us |
| L1 | CG | 22A | 28 peptide + 200 DMPC + 1 open LCAT in 20x20x20 nm<sup>3</sup> | 40 us x3 |
| L2 | CG | 22A-R7Q | " | 40 us x3 |
| R1 | CG | 22A-R7D | " | 20 us |
| R2 | CG | 22A-R7H | " | 20 us |
| R3 | CG | 22A-R7C | " | 20 us |
| R4 | CG | 22A-R7L | " | 20 us |
| R5 | CG | 22A-R7K | " | 20 us |
| I1 | AA | 22A | 9 peptide + 60 DMPC + 1 open LCAT in 12x12x12 nm<sup>3</sup> | 600 ns + 300 ns x2 |
| I2 | AA | 22A-R7Q | " | 300 ns x3 |
| I3 | AA | apoA1 helix 4 | " | 300 ns |
| I4 | AA | apoA1 helix 5 | " | 300 ns |
| I5 | AA | apoA1 helix 6 | " | 300 ns |
| I6 | AA | apoE helix 1 | " | 300 ns x2 |
| I7 | AA | apoE helix 6 | " | cancelled |
| M1 | CG | 22A-D4N | 28 peptide + 200 DMPC + 1 open LCAT in 20x20x20 nm<sup>3</sup> | 20 us |
| M2 | CG | 22A-D4N-R7Q | " | 20 us |
| N1 | CG | apoA1 helix H | " | 20 us |
| N2 | CG | apoA1 helix I | " | 20 us |
| N3 | CG | apoA1 helix I kinked | " | 20 us |

**Table 1** continued
| ID | res | peptide | composition | runtime |
| --- | --- | --- | --- | --- |
| N4 | CG | apoE helix B | " | 8 us |
| N5 | CG | apoE helix C | " | cancelled |
| N6 | CG | apoE helix H | " | 20 us |
| N7 | CG | apoE helix J | " | 8 us |
| N8 | CG | apoE helix L | " | 8 us |
| N9 | CG | apoB100 helix M | " | cancelled |
| N10 | CG | apoB100 helix X | " | 8 us |
| N11 | CG | apoB100 helix Z | " | cancelled |
| N12 | CG | apoB100 helix AB | " | 8 us |
| N13 | CG | apoB100 helix AD | " | cancelled |
| N14 | CG | apoB100 helix AE | " | 8 us |
| N15 | CG | apoB100 helix AF | " | 8 us |
| N16 | CG | apoB100 helix AG | " | 8 us |
| N17 | CG | apoB100 helix AN | " | 8 us |
| N18 | CG | apoB100 helix AO | " | 8 us |
| N19 | CG | apoB100 helix AP | " | 8 us |
| N20 | CG | apoB100 helix AQ | " | 8 us |
| N21 | CG | apoB100 helix AS | " | 20 us |
| N22 | CG | apoC1 helix A | " | 8 us |
| N23 | CG | apoL1 helix D | " | 20 us |
| N24 | CG | apoL1 helix E | " | 8 us |
| N25 | CG | LCAT helix D | " | 8 us |
| N26 | CG | serum amyloid A1/2 helix B | " | 20 us |
| N27 | CG | albumin helix I | " | cancelled |
| B1 | AA | apoB100 helix AS | 9 peptide + 60 DMPC + 1 open LCAT in 12x12x12 nm<sup>3</sup> | 300 ns |

### Apolipoprotein fragment screen finds LCAT-binding helixes

In CG MD simulations with LCAT in its open conformation and a nanodisc composed of apoA1 mimetic peptides and DMPC the peptides bind to the lid cavity of LCAT [15]. We hypothesize that the helixes of apoA1, which the mimetic peptide 22A is based on, utilize the same site. Each helix of apoA1 and apoE were screened by simulating them with a lipid bilayer and open LCAT (A1-8 and E1-8 systems). If a peptide was not able to attach to the lipid membrane at all due to high hydrophilicity or lack of amphiphilicity, the simulation was cancelled, as we are interested in helixes that favour both the lipid and LCAT environment. An overview of the system and binding site is shown in Figure 1A-B. Besides occupancy, which is the percentage of time LCAT was occupied by a peptide, we also measured the peptide-peptide dimerization tendency and configuration (Figure 1C) and the percentage of peptides dissociated from the lipid bilayer. The results of the screen are shown in Figure 1D-E and Table 2.

**Figure 1.**
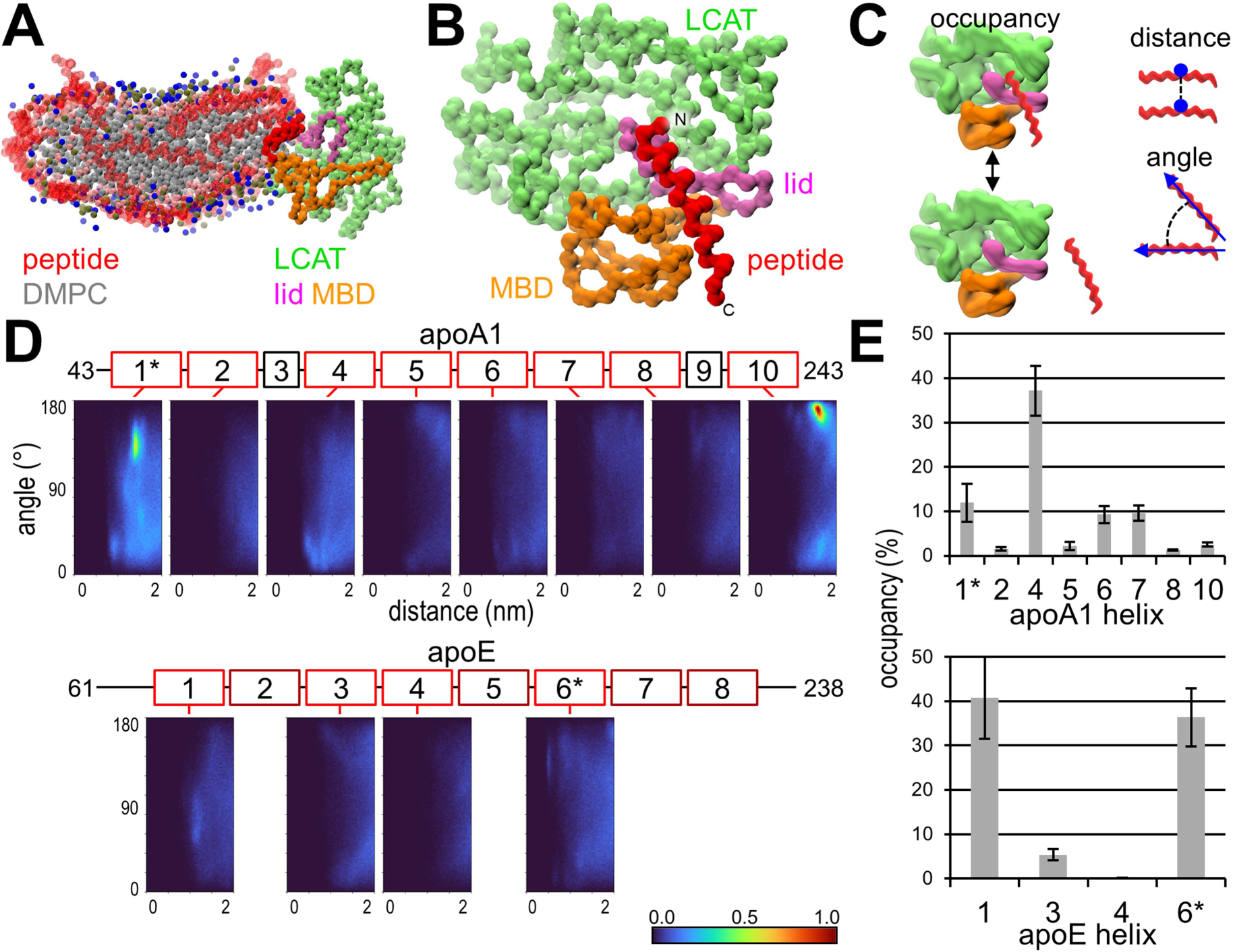
Coarse-grained systems’ characteristics and apolipoprotein results. A) A snapshot of the 22A simulation showing the position LCAT takes in the perimeter of the nanodisc. Peptides are transparent apart from the bound peptide. B) The peptide binding site on LCAT when LCAT is on the nanodisc. Rest of the nanodisc is not shown. C) Illustration of the primary parameters measured with the system. D) Angle-distance dimerization profiles for the apolipoprotein helixes. Apolipoprotein sequences were split by their helical assignment and simulated separately as helical peptides. The colorbar is normalized by the maximum bin count of apoA1 helix 10. E) Occupancy results for the apolipoprotein helixes. *secondary structure predicted by PEP-FOLD3 wasn’t helical, but simulated in a helical conformation

**Table 2.** Apolipoprotein fragment screen results.

| apolipoprotein | helix ID | residues | occupancy (%) | dissociated (%) | hydrophilicity |
| --- | --- | --- | --- | --- | --- |
| apoA1 | 1* | 44-65 | 11.9±4.3 | 18.3±0.2 | 0.1 |
|  | 2 | 66-87 | 1.6±0.4 | 30.6±2.6 | 0.54 |
|  | 4 | 99-120 | 37.2±5.6 | 16.8±1.6 | 0.62 |
|  | 5 | 121-142 | 2.2±0.9 | 53.8±0.8 | 0.65 |
|  | 6 | 143-164 | 9.3±1.9 | 39.7±1.2 | 0.56 |
|  | 7 | 165-186 | 9.6±1.7 | 49.6±1.9 | 0.62 |
|  | 8 | 187-208 | 1.3±0.2 | 46.2±1.2 | 0.44 |
|  | 10 | 220-243 | 2.5±0.5 | 1.3±1.1 | 0.02 |
| apoE | 1 | 62-83 | 40.8±9.3 | 62.7±1.1 | 0.62 |
|  | 2† | 84-105 | - | - | 0.54 |
|  | 3 | 106-127 | 5.3±1.2 | 38.7±1.2 | 0.15 |
|  | 4 | 128-149 | 0.0±0.1 | 47.6±0.6 | 0.64 |
|  | 5† | 150-171 | - | - | 0.80 |
|  | 6* | 172-193 | 36.4±6.5 | 30.8±4.9 | 0.45 |
|  | 7† | 194-215 | - | - | 0.17 |
|  | 8† | 216-237 | - | - | 1.12 |

\*secondary structure predicted by PEP-FOLD3 wasn't helical, but simulated in a helical conformation
†cancelled due to practically 100 % dissociation

We propose that the significance of the helix binding site is to facilitate lipid entry by either improving the positioning of LCAT respective to the bilayer and or by locking the lid into a favourable configuration. Thus, the LCAT-nanodisc interaction takes the following steps. First, LCAT binds to the nanodisc with the MBD, as the kinetics of this initial attachment have been shown to be equal between different lipoprotein particles [37]. Second, LCAT, with its closed lid protecting the hydrophobic active site from the aqueous environment, migrates to the perimeter of the disc, which was seen in CG MD simulations of closed LCAT and was further supported by EM imaging, although EM couldn’t confirm the state of the lid [2,15]. Third, the lid of LCAT opens due to entropic forces and likely through the help of the lipid or peptide components of the nanodisc. Fourth, the stabilizing peptides position the active site opening and keep the lid open for cholesterol and phospholipids to enter and the resulting CE and lysolipid to exit. Finally, after one catalytic cycle LCAT detaches and either during this process or after the lid is driven back to the closed state [38].

For apoA1, helixes 4, 6 and 7 had clearly higher binding to LCAT than the other helixes. Helix 6 has been identified as a likely interaction site for LCAT based on hydrogen exchange, cross-link, and QCM studies as well as discovered mutations [2,30,39,40]. These studies are evaluated in more detail with our atomistic simulations. Relative to helix 4 and 22A, helixes 6 and 7 presented with a high dissociation (∼40-50 % vs ∼0-15 %) and low occupancy (∼10 % vs ∼35 %). Our reasoning is that although separate helixes are more hydrophilic than 22A, when they are linked together in apoA1 the increased mass decreases their hydrophilicity enabling them to remain attached to the lipid bilayer. The higher mass also reduces movement, which may otherwise lead to LCAT detachment. Regardless, a later screen identified a more preferable binding site between helixes 6 and 7(N2 system), which we will discuss later. For apoE, half of its helixes were too hydrophilic to form nanodiscs, but out of the remaining four 1 and 6 had an occupancy similar to helix 4 of apoA1 and 22A.

The calculated average hydrophilicity values showed that the helixes at the end of apoA1 had a low hydrophilicity, while for middle helixes it was high. This was reflected moderately well in the fraction of helixes that were dissociated from the lipid bilayer (R^2^=0.53 with simple linear regression). We later explored this trend with all non-22A based CG systems. If cancelled systems were taken to consideration with dissociation set to 100 %, dissociation vs hydrophilicity resulted in R^2^ of 0.28 (positive correlation) whereas dissociation vs hydrophobic moment resulted in R^2^ of 0.37 (negative correlation). If cancelled systems were left out (i.e. only accurate dissociation values were used) the R^2^ values were 0.38 and 0.24 respectively. The dissociation of apoA1 helixes strongly correlate with their experimental logarithmic partition coefficients attained from a lipid binding assay (R^2^=0.75) [41].

To test whether full length apoA1 would have improved binding kinetics as proposed, we simulated a small nanodisc with a cyclical apoA1 (S1 system). This system was a product of earlier unsuccessful pilot systems made to sample LCAT binding to apoA1. By placing helix 4 to the lid cavity LCAT was able to remain attached to helix 4 for ∼8 µs (Figure S1). This is remarkably high considering that the longest binding event for 22A lasted ∼200 ns. Besides the reasons given above, the stronger interaction with LCAT may have been caused by the additional interactions with helix 3 and by the missing N-terminal charge. Thus, it is possible that some of the cancelled helixes of apoE which didn’t bind to the lipid bilayer would perform well in the context of the whole protein. As this system requires more assumptions on the secondary structure of apoA1, we were satisfied with this qualitative result.

The secondary structures of all screened peptides were evaluated with PEP-FOLD-3 [35], which revealed that the conformation of helix 1 wasn’t completely helical and confirmed that the proline in apoE helix 6 induces a kink. This is a limitation of the CG models we defined, which force the peptides to a helical conformation. The conformation could be amended by reducing helicity or by introducing a kink, but we decided against this strategy as our unpublished simulations indicate that increased conformational freedom reduces occupancy. We later explored the effect of a kink with a more promising peptide (N3 system) and will return to apoE helix 6 results in that context. Although peptide helicity increases when it’s bound to a lipid bilayer [42], we will consider these helixes as poor binding for now.

LCAT independent angle-distance dimerization plots were generated for all peptide pairs. The unnormalized profiles are in Figure S2. Helix 10 of apoA1 has a clear antiparallel dimer hotspot and helix 5 presented with a faint 22a-like profile. Curiously, reconstituted HDL presented with no intra- or intermolecular crosslinks between helix 10s, but human HDL did [43]. The profile of helix 5 supports the antiparallel double belt model of apoA1 with a 5/5 registry [1]. Although the dimerization was faint compared to helix 10, the sampling was likely harmed by the high dissociation from the lipid membrane. In 1:1 mixed systems of apoA1 helixes 4 and 6 no clear dimers formed between them (S2 system), although a slight preference for antiparallel dimerization was present.

### Acids of residue 4 and arginine of residue 7 anchors peptides to LCAT

Having identified the best binding helixes of apoA1 and apoE, we investigated which residues remained similar between helixes 4, 6 and 7 of apoA1, while being different to apoE helix 1. Particularly the first halves of the helixes were looked at, which have the most preference to the binding site of LCAT. The residues at positions 7 and 9 stood out. In apoA1 helixes position 7 is Q, R and R respectively and position 9 is K, R and R, whereas for apoE helix 1 they are M and E. To confirm which residues are the most favourable for LCAT binding, we screened some R7 mutants of 22A (R1-5 systems). The results are shown in Figure 2, along with 22A and 22A-R7Q (L1-2 systems). This screen afforded us two insights. First, for 22A, which is based on apoA1, neutral or positively charged residues at position 7 with affinity for hydrogen bonding seem to perform well, but overall R and Q perform best. Second, the very strong antiparallel dimerization induced by R7C and R7L presumably harms the peptides’ ability to occupy LCAT.

**Figure 2.**
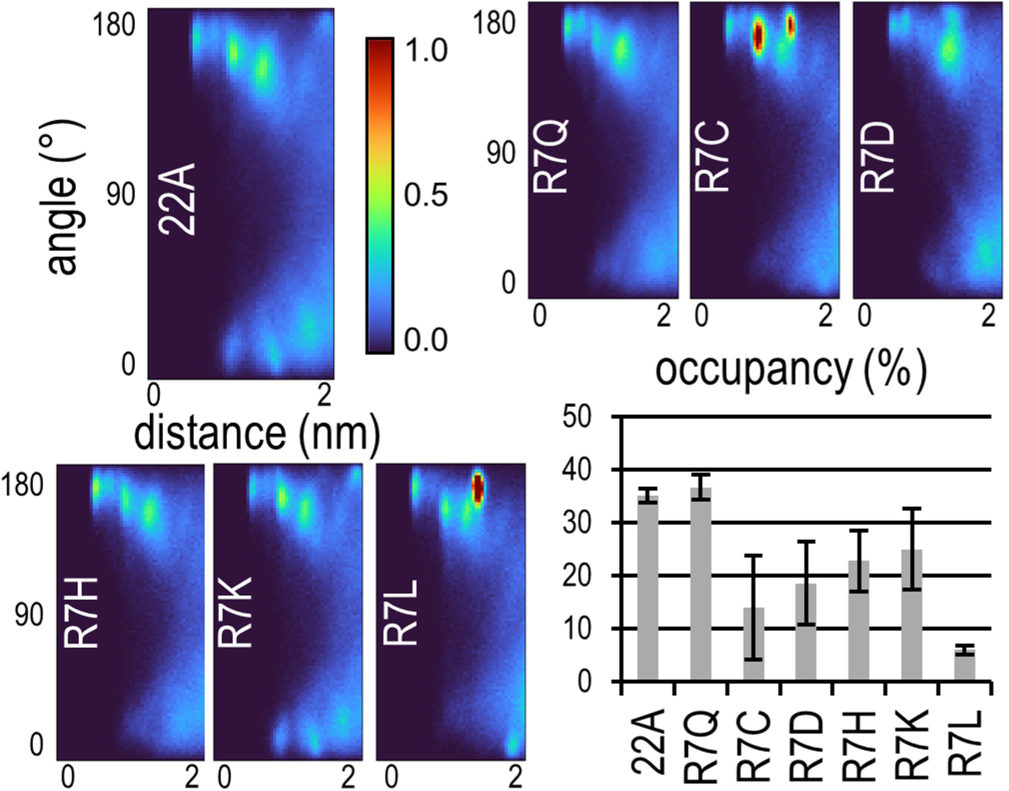
R7 mutant screen results. The colorbar of angle-distance dimerization profiles is normalized by the maximum bin count of apoA1 helix 10 shown in Fig 1. The peaks in R7C and R7L are 34 % and 147 % above the scale respectively.

Given that arginine is capable of forming salt bridges, which we suspect to make the key interactions between LCAT and peptides in the binding site, it is surprising that neutralizing it to glutamine doesn’t harm occupancy [15]. Thus, we decided to explore the binding interactions of 22A and 22A-R7Q in atomistic resolution by preparing smaller nanodisc systems and atomizing them when a peptide occupies the binding site (I1-2 systems). In all three simulations R7 of 22A indeed forms a salt bridge to D73 of LCAT and in one simulation D4 of 22A formed hydrogen bonds with S236 of LCAT. These were the only stable hydrogen bonds between LCAT and the peptide. During the simulations 22A shifted away from the binding site slightly, pointing away from LCAT, while maintaining the R7-D73 salt bridge (Figure 3A-B).

**Figure 3.**
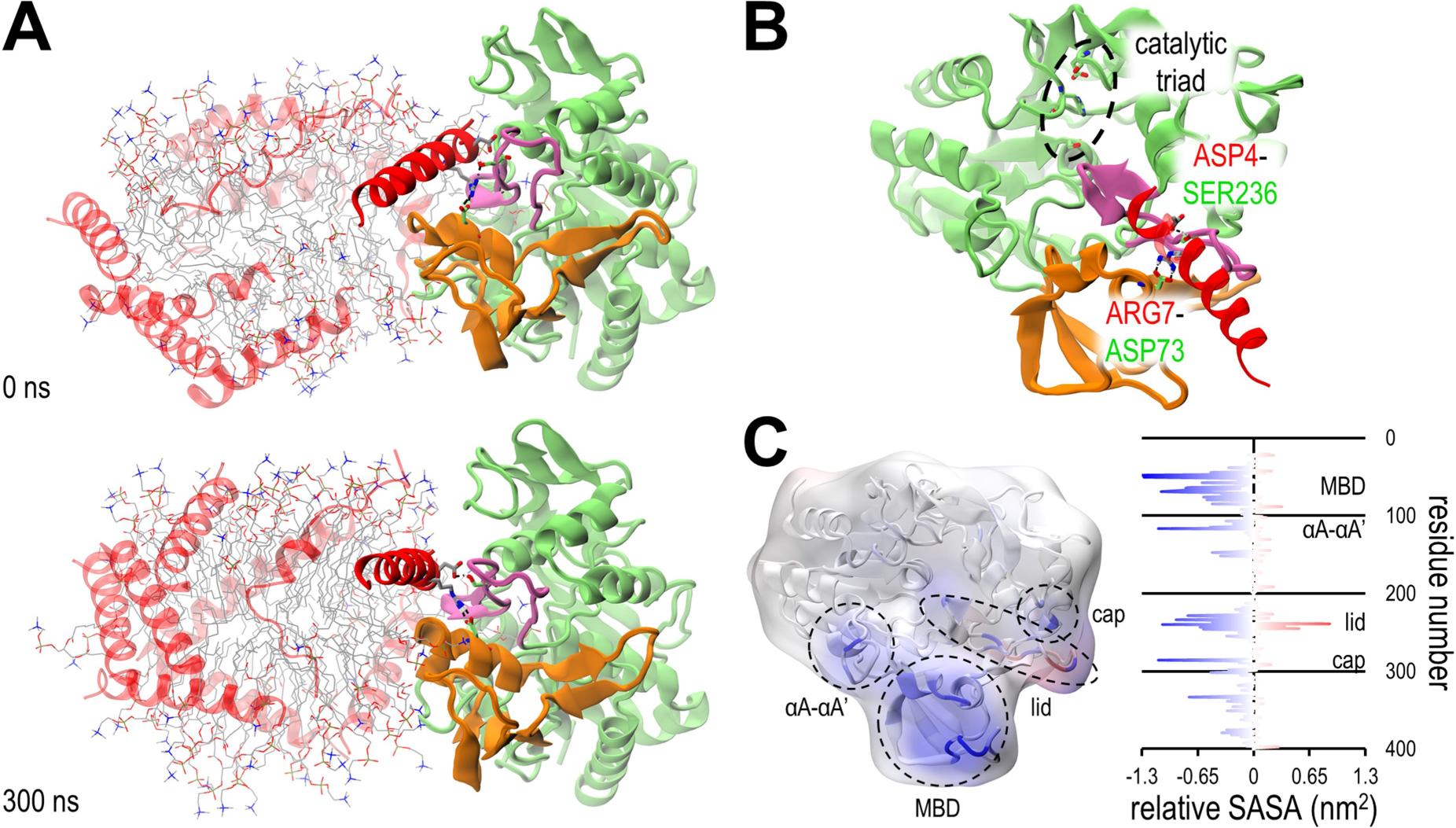
The LCAT-nanodisc interaction. A) Snapshots of the start and end configurations of an atomistic simulation with 22A based nanodisc and LCAT. Colouring follows the conventions of Figure 1. LCAT and peptides rendered as cartoon, relevant residues as sticks, hydrogen bonds as dashed lines and DMPC as lines. B) The peptide binding site of the snapshot at 300 ns shown in more detail. Rest of the nanodisc is not shown. Although the D4-S236 bond didn’t occur in all 22A simulations, it is shown here as other peptides commonly utilized it. C) Relative solvent accessible surface area per residue between closed LCAT and open LCAT with nanodisc. The results are mapped onto a cartoon rendering and surface map of LCAT superimposed. The negative values marked with blue indicate residues that are protected from water when LCAT binds to a nanodisc. Red indicates less protection from water.

If the bound 22A was instead a helix of apoA1, this new position would enable a preceding helix to fit over the active site of LCAT and the other apoA1 of the double belt model to fit on top of the bound apoA1. In this position, cholesterol and phospholipids aiming to enter LCAT would interact with LCAT’s αA-αA’ loop (residues 112-120), top loop of the MBD (61-75) and lid loop (227-247), as well as with the bound and preceding helixes of the double belt. We had previously compared existing hydrogen-deuterium exchange (HDX) mass spectrometry results to CG simulation derived per residue solvent accessible surface areas (SASA) [2,15]. In the HDX experiment the HDX difference between of LCAT in solvent and LCAT with reconstituted HDL (carried out with apoA1) was measured, which could be roughly replicated in silico by measuring the SASA difference of closed LCAT in water and open LCAT on nanodisc. Performing this analysis with the atomistic systems revealed that, as expected, the atomistic resolution offered a better fit HDX data (Figure 3C). The fit isn’t perfect, particularly in the cap domain the simulations have less protection than the HDX results would suggest, but as stated the HDX data is measured for apoA1 based HDL, which may position LCAT slightly differently than 22A based nanodiscs. Furthermore, the short timescales the simulations offer results in insufficient sampling of e.g. the many conformations of the lid that are captured in the HDX results. In all the slow HDX regions the SASA analysis identified residues with increased protection, which supports the presented binding model. The binding model also doesn’t occlude the glycosylation sites or the known binding site of the agonistic 27C3 antibody [44].

Breaking the R7-D73 salt bridge through 22A-R7Q resulted in a bond between D4 and S236 of LCAT in two simulations and in a salt bridge between D4 and K238 in one simulation. 22A-R7Q took the same position as 22A. Given that in one of the 22A simulations the D4-S236 bond appeared, it is likely that both 22A and 22A-R7Q bind to LCAT through D4 and 22A additionally utilizes R7. As their induced V_max_ values are similar the R7-D73 salt bridge seems supplementary. The atomistic resolution additionally revealed that unbound 22A maintains a salt bridge/hydrogen bond between its D4 and R7, which is missing from the LCAT bound peptide and from all R7Q mutants. To confirm that the CG model was using D4 for binding we screened 22A-D4N and 22A-D4N-R7Q (M1-2 systems), resulting in occupancies of 5.6±0.4 % and 2.8±1.3 % respectively.

The bonding capability of D4 prompted us to return to the apolipoprotein helix sequences. For apoA1, only the well binding helixes 4, 6 and 7 had a combination with D or E at position 4 and R or Q at position 7. We will call this combination the DE4-QR7 moiety. Helix 10 had an E4-K7 configuration, which, like 22A-R7K, performed worse than R7 or Q7 combinations in the CG MD binding screen. For apoE, only helixes 2, 4 and 5 had the DE4-QR7 moiety, where helixes 2 and 5 didn’t bind to the lipid bilayer sufficiently and helix 4 had no affinity for LCAT. The well binding helixes 1 and 6 had D4-M7 and S4-R7 combinations instead. Notably, out of all peptides we have screened only apoE helix 6 presented with a capacity to bind to LCAT without having D or E at position 4, although, as discussed above, it was forced into an unnatural secondary structure.

As the best helixes of apoA1 had the DE4-QR7 moiety, whereas the helixes of apoE didn’t, we simulated them in an atomistic resolution like 22A and 22A-R7Q (I3-7). Simulations of apoE helix 6 were attempted, but helix 6 rapidly unfolded into different conformations. Overall, the attained data from the atomistic simulations is limited by insufficient sampling. As both the peptide and the random chain lid of LCAT can take a multitude of positions respective to each other, a lot of long lasting binding configurations can form with heterogeneous interactions. Proper sampling would be attained by running several systems in parallel, but the large size of the systems limited us. Therefore, we will only highlight the interactions we are most confident in.

Helixes 6 and 7 of apoA1 both bound to LCAT through R7-D73 and the E4 of helix 6 bound to K238 whereas the D4 of helix 7 bound to S236. Helix 4 of apoA1 which lacks R7 instead utilized K8-D73 and D4-S236. Helix 1 of apoE was run in duplicate. In one simulation it bound to D73 through K11 whereas in the other it formed short bonds to S236 through D4. Treating the apoA1 results in bulk shows that apoA1 based peptides primarily bind to LCAT via D4-S236 or D4-K238 and if R7 is available via R7-D73. Helix 1 of apoE likely has the same capability to bind to S236 and K238 through D4, but it might utilize additional interactions with K11. It is possible helixes 2 or 5 of apoE, which have the DE4-QR7 moiety, could bind with apoA1-like interactions to LCAT, but due to the poor lipid binding this couldn’t be evaluated.

The equilibrated binding pose the peptides take in atomistic simulations allows us to estimate how the binding model fits with in vitro LCAT-apoA1 crosslink and EM data [1,2]. The binding of helixes 4, 6 or 7 explain all of the crosslinks apart from ones involving LCAT K159 or S255, as those are located behind LCAT relative to the lipid bilayer (Figure 4A). In fact, LCAT K159 is part of an amphiphilic helix which we later discovered to be capable of forming nanodiscs in silico (LCAT helix D). Thus, for our binding model to be valid the K159 crosslinks must be caused by random tumbling and perhaps partly through helix-helix interactions between backwards LCAT and apoA1. Overall, both simulations and crosslinks agree that LCAT has many potential binding sites on apoA1. Helix 4 binding explained four crosslinks, helix 6 nine and helix 7 four. Helix 7 explained the most common BS3 based crosslink and helix 6 the two next most common ones. If an extra 6 Å is added to the distance requirement due to dynamics helix 4 also explains these latter two crosslinks. Out of the four highest scoring DC4 based crosslinks helix 4 explains one, helix 6 three and helix 7 three.

**Figure 4.**
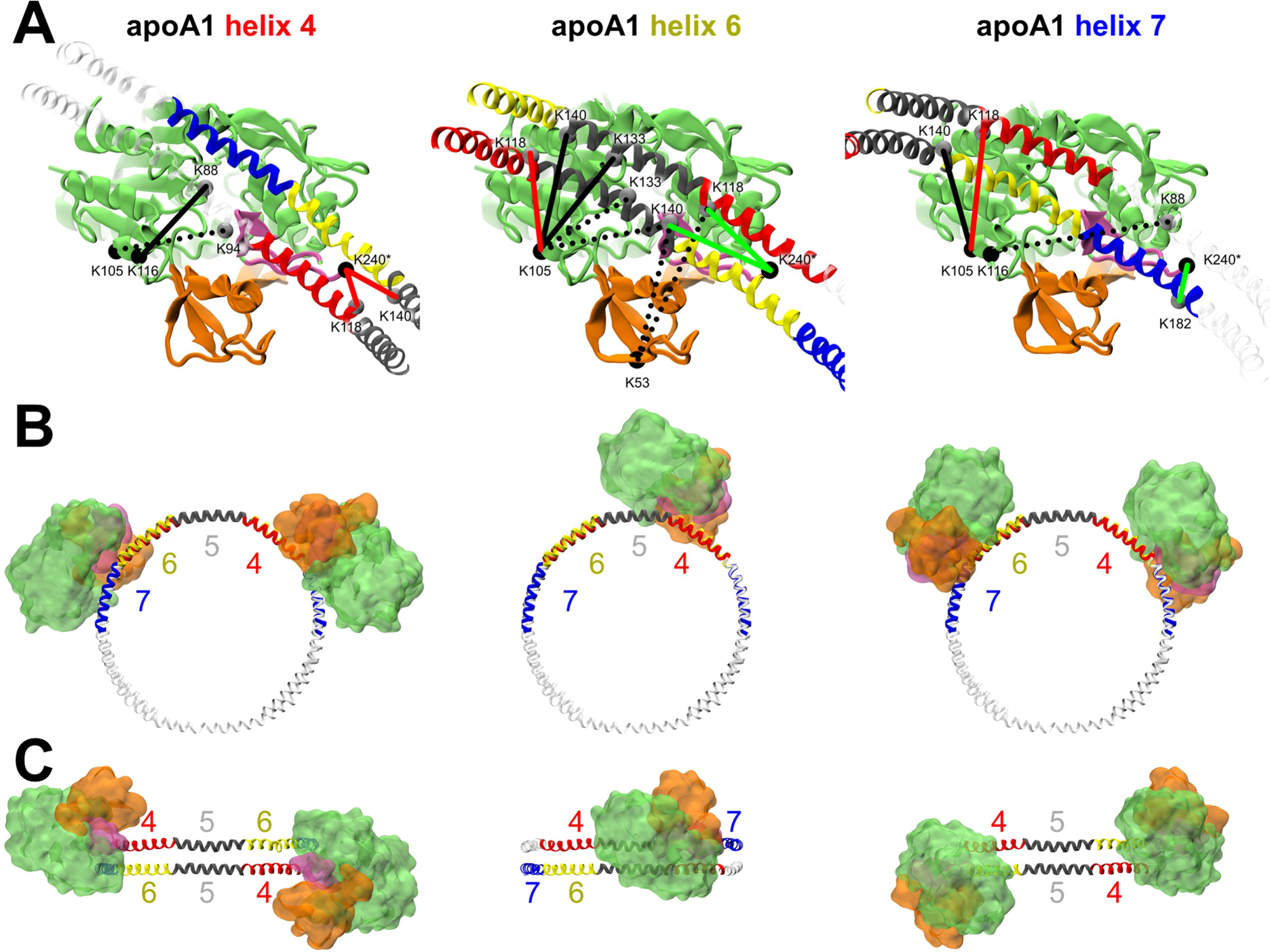
Different binding spots LCAT would take on nHDL with the proposed binding model explored. A: Known cross-link pairs whose Cα-Cα distance is within 40 Å are indicated by lines. Green and red lines indicate BS3 cross-links whose distance requirement of 24 Å is met or not met respectively. Dashed lines indicate links with significant occlusion. B: What nHDL would look like from the perspective of the disc normal if two LCATs were bound to either helixes 4, 6 or 7 of apoA1. For helix 6 only one is shown due to occlusion. C: Same visualization from the perspective of the disc perimeter.

The EM data for apoA1 in the 5/5 registry with two LCATs bound only supports helix 4 in this binding model (Figure 4B-C) [2]. If LCAT was bound to helixes 6 or 7, the lobes representing LCAT’s density would be positioned on the opposite sides respective to the disc normal. However, binding to helixes 4 still leaves the angle between LCATs in the upper range (∼90 degrees). The 5/4 registry would decrease the angle [43], as well as LCAT utilizing other DE4-QR7 moieties in helix 4, e.g. E113-R116 or E120-R123, closer to the experimentally measured 63±10 degrees. Although not supported by the EM density map, as ∼1 % of particles presented with three or more LCATs bound, LCAT may also utilize several helixes of the same apoA1. Simultaneous binding to helixes 4 and 6 of the same apoA1 would produce an angle closer to the detected distribution. Another factor supporting unspecific binding is that on nanodisc particles made of 22A LCAT also has a tendency to bind ∼60 degrees apart, which suggests that another mechanism, such as the dimerization of LCAT, is behind the angle distribution [15]. Dimerization would explain LCAT-LCAT K105 crosslinks that otherwise would require simultaneous helix 6 and 7 binding in 5/5 registry or helix 6 binding in 5/4 registry with an adjusted LCAT orientation in this binding model [2].

Curiously, although binding to helix 4 seems to satisfy some crosslink and EM results relatively well, HDL-C harming mutations have been largely detected in helix 6, while mutations in helix 4 maintain HDL-C [39]. An R7Q mutation in helix 6, i.e. R149Q, harms LCAT acyltransferase activity markedly [45]. In fact, helix 6 contains three DE4-QR7 moieties: E146-R149, D150-R153 and D157-R160, and when any of these arginines are mutated to glutamine V_max_ drops to ∼10 %. Conversely, if a charge is removed from any of the acids, V_max_ drops to ∼50 %. From the limited view the atomistic simulations afforded us, there are no evident interactions at play that would explain this activity loss if apoA1 is attached to the peptide binding site of LCAT via E146-R149. Instead, it may arise from a loss of helicity caused be a disturbance in α-helix stabilizing i, i+3 type salt bridges [46]. If LCAT was bound to helix 7, helix 6 would be interacting with lipids entering the active site of LCAT and thus lipid entry would be sensitive to the structure and nature of helix 6. This would explain why losing any of the helix maintaining salt bridges causes such a loss in activity, as well as the many HDL-C harming mutants. In apoA1 helix 7, much like in 22A, an R7Q mutation (R171Q) didn’t harm activity [45].

Cooke et al. collected a list of apoA1 mutants with decreased LCAT activity without decreased ABCA1 cholesterol efflux capabilities [1]. Helix 6 R7 mutant R149V results in a V_max_ of 32 % compared to wild type, likely through a similar mechanism as above [47]. Disrupting the D4 of helix 4 through a D102A+D103A mutant drops V_max_ to 70 % whereas a D168A at D4 of helix 7 drops it to 38 % [48,49]. Similarly, K_m_ rose to 136 % and 300 % respectively. The helix 6 mutations instead generally improved or didn’t affect K_m_ apart from R160Q (164 %) and E146A+R149A (131 %) [45]. The mutation data supports several binding sites, but any activity losses from these mutants can’t be purely attributed to broken LCAT-apoA1 interactions since they can’t be decoupled from α-helix or apoA1-apoA1 stabilizing salt bridges.

No LCAT mutations that directly affect the interactions of interest were found. The search did afford 4 mutations that plausibly interact with apoA1 when it’s bound to the LCAT binding site. FED causing V46E [50] and FLD causing D77N [51], N228K [52] and G230R [53]. Given that the lid has a relaxed conformation and several functions, such as modulating LCAT activity via uncovering the active site, potentially maintaining lipid entry and exit, and binding to lipoprotein particles via interactions to lipids and apolipoproteins, lid mutations (N228K and G230R) can affect several of these pathways. Thus, out of this group, the MBD mutants V46E and D77N are more likely to cleanly affect only the LCAT-apolipoprotein interaction.

### The DE4-QR7 moiety revealed LCAT binding fragments in other plasma proteins

The evidence presented so far has shown that the presence of a DE4-QR7 moiety has some power in predicting a helical peptide’s capability to bind to LCAT. We decided to attempt finding novel protein fragments that could bind to LCAT with the same mechanism in the context of a lipid bilayer (N1-27 systems). The sequences of apolipoproteins A1, E, C1, C3, B100, D, F, H, L1, M and (A) as well as the sequences of LCAT, serum amyloid A (SAA) isoforms 1, 2 and 4, albumin and prothrombin were screened to find 22 amino acid long fragments that have a D or E at position 4 and Q or R at position 7. The sequences of apoA1 and apoE were scrutinized again to identify all DE4-QR7 containing fragments that may be outside the standard helix categorization of residues. Each detected fragment was given a letter identifier in order of appearance (Table S2). Mean hydrophilicities and helical penalties were calculated for each fragment, and ones that had values lower than 0.75 and 2 kJ/mol respectively were put through PEP-FOLD3 to assess their conformation. Suitably helical fragments were screened with the standard CG MD LCAT system. An exception in the helicity requirement was made for helixes H and I of apoA1, which contain the other two moieties of helix 6. All systems were run to 8 us, and continued to 20 us if they had good binding. The occupancy results are shown in Table 3 and the unnormalized angle-distance dimerization profiles for all systems in Figure S2. Dissociation was also noted to affect LCAT positioning (Figure S3).

**Table 3.**
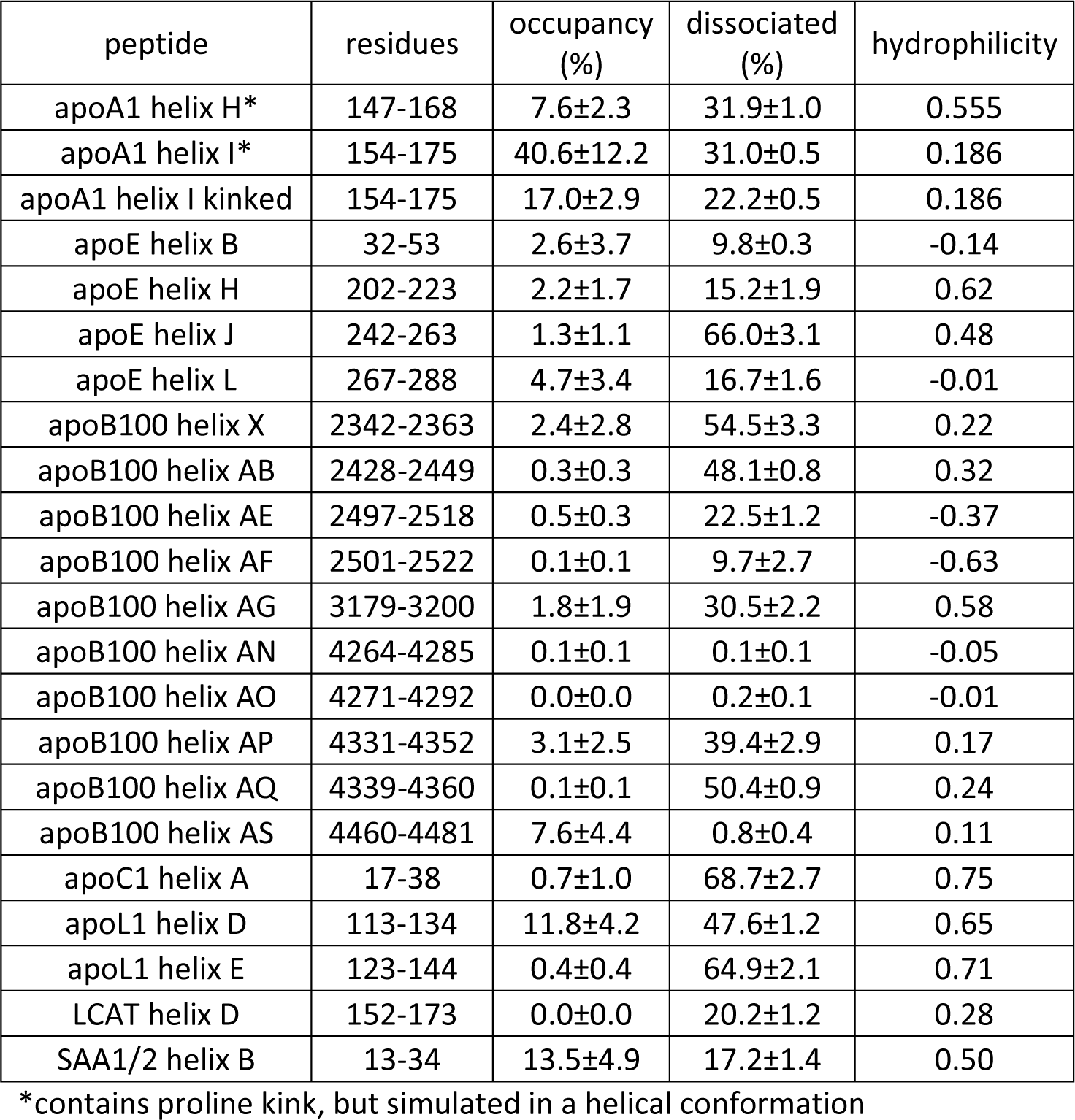
Plasma protein screen results.

Both apoA1 helix 6-7 based peptides presented with an ability to bind to LCAT. Helix I, located at residues 154-175, had an occupancy of ∼40 %, which is in line with the values of helix 4 and 22A as well as the best binding helixes in apoE. However, as discussed with apoE helix 6, the true binding tendency is hard to evaluate as the conformational freedom the proline kink introduces is missing from the CG model. As in vitro data strongly indicated that the helix 6-7 region of apoA1 is a binding site of LCAT we decided to simulate helix I with a kinked conformation. The kink was set to the peptide’s proline at position 12, which, curiously, is in the same position as the proline of apoE helix 6. This still afforded a relatively high occupancy of 17.0±2.9 %, although the value is likely overestimated since the condition for occupancy was defined for purely helical peptides. Still, taking into consideration that the surrounding apolipoprotein chains prevent larger kink rotations, the high occupancy suggests that apoA1 helix I and apoE helix 6 are valid binding sites for LCAT.

Four other systems were continued to 20 µs: apoE helix H, apoB100 helix AS, apoL1 helix D and SAA1 helix B. Out of these the latter three had a favourable binding profile. The residues of helix D are located at positions 113-134 (numbering without signal peptide) in the pore forming domain of apoL1. This part of the domain is predicted to locate in the extracellular space [54]. Interestingly, the tail of helix D also contains the start of the BH3 domain responsible for regulating apoptosis [55]. However, although PEP-FOLD3 predicted a helical conformation for helix D, in crystal structures of apoL1 the proline at position 6 introduces a kink (PDB IDs: 7LFA, 7LFB). Thus, it is unlikely LCAT would interact with the group as predicted. Helix E, which is located at positions 123-144 and contains the BH3 domain completely, had no affinity for LCAT.

ApoC1 has known LCAT activity [56]. Notably, a peptide fragment of residues 24-57 induced a 60 % activity in LCAT, whereas a fragment with residues 17-57 induced 100 %. This latter fragment included the DE4-QR7 moiety starting at position 17, which was also the starting residue of helix A, which surprisingly didn’t bind to LCAT in the CG MD binding screen. A potential reason is that NMR studies have identified a flexible hinge at positions 30-37 so LCAT may employ a different binding technique instead [57]. Starting at position 41 another DE4-QR7 moiety is present but this wasn’t tested as it would form only a 17 residue long peptide. A fragment of residues 39-57 couldn’t activate LCAT, but neither could it form stable complexes, which may have impeded LCAT binding [56].

SAA proteins are associated with HDL. They are capable of displacing apoA1 to form dysfunctional HDL with decreased cholesterol efflux [58]. Interestingly, helix B of SAA1 and SAA2 (identical sequence), which is located at positions 13-34, has a relatively high affinity for LCAT. The crystal structures of SAA1 do include a flexible hinge at ∼15 residues into the helix (PDB IDs: 4IP8, 4IP9), but if the LCAT binding part of the helix is stable it may not harm binding. This suggests that SAA rich HDL may also be partly dysfunctional due to reduced acyltransferase activity caused by apoA1 competing with SAA for LCAT binding.

One LCAT binding fragment was detected in apoB100: helix AS, which is located at residues 4460-4481. Helix AS along with helix AN and apoE helix H were the only ones with a strong tendency to form antiparallel dimers out of the N1-27 systems (Figure S2). Knowing that this was the first LCAT binding site directly associated with LDL, we simulated the helix in atomistic resolution using the system described above for 300 ns (B1 system). Much like apoA1-like peptides apoB100 helix AS used D4 to bind to S236 of LCAT. Lacking R7, helix AS instead used K10 to salt bridge to D73 of LCAT, similarly to how apoE helix 1 used K11. Once again, only the most relevant interactions are highlighted due to limited sampling.

It is worth repeating that the CG MD binding screen utilized here forces the peptides into a helical conformation and is only usable if they have some tendency to complex with lipids. Furthermore, coarse-graining naturally loses some detail on interactions between functional groups such as the directionality of hydrogen bonds. A testament to this is the slightly different binding pose the peptides take on LCAT in atomistic resolution. Thus, it is very likely that the screens missed some good binding peptides or highlighted peptides that bind poorly. Taking the atomistic data together, perhaps a better predictor for LCAT binding would be a DE4-R7/KR10/KR11 moiety. Regardless, it is evident that the validation of the presented model requires in vitro screens with mutated LCAT, and especially mutants of D73, S236 and K238 should be considered. In terms of selectively inhibiting LCAT β-activity, the discovered binding site and interactions pave the way to maintain LCAT apoA1 interactions and diminish apoE or apoB100 interactions. Given that the same residues of LCAT are targeted for hydrogen bonding and salt bridging, more subtle ways of sterically hindering bulky side chains or reducing hydrophobic contacts may be more fruitful.

## CONCLUSION

CG and atomistic MD simulations were utilized to characterize the LCAT-nanodisc and LCAT-HDL interactions in higher detail. Through a CG MD screen it was discovered that the helixes of apoA1 are able to utilize the same binding site on LCAT as apoA1 mimetic peptides. The screen identified helixes 4, 6 and 7 as well binding, which is supported by in vitro studies of previous literature. Atomistic simulations further revealed that the binding determinants that drive the attraction between apoA1-like helixes and LCAT are the formation of hydrogen bonds or salt bridges between peptide E4 or D4 and LCAT S236 or K238 residues and salt bridges between peptide R7 and LCAT D73 if R7 is available. We employed this knowledge to screen for novel LCAT binding helixes in different plasma proteins. Such helixes were detected in apoL1, apoB100 and serum amyloid A2. Our results highlight that LCAT β-activity on LDL, where apoE and potentially apoB100 are hypothesized to interact with LCAT, may be driven by the same E4 or D4 -S236 and R7-D73 binding determinants, except the missing R7 is replaced by lysines at positions 10 or 11. The findings support the development of therapeutic nanodiscs based on apoA1-mimetic peptides in the context of LCAT deficiencies and cardiovascular diseases. The modulation of α- and β-activities paves the way for novel LCAT biologics and for facilitating the investigation of the distinct roles of these activities in cardiovascular disease.

