## Supplementary Information for "Systematic Evaluation of Lecithin:Cholesterol Acyltransferase Binding Sites in Apolipoproteins via Peptide Based Nanodiscs: Regulatory Role of Charged Residues at Positions 4 and 7"

**Table S1.** Peptide sequences in simulated systems.

**Table S2.** All 22 residue long sequences with DE4-QR7 moieties in a variety of proteins.

**Figure S1.** LCAT binding pose from different perspectives on small nanodisc with one cyclic truncated apoA1. Residues 44-241 of apoA1 were used in the model. LCAT is coloured according to Figure 1 and apoA1 helixes according to Figure 4.

**Figure S2.** Angle-distance profiles of CG systems with unnormalized colorbars. The S2 system's bin counts are not comparable to other systems as it had 14 x 14 peptide pairs, unlike the other systems with 378 peptide pairs.

**Figure S3.** LCAT position density plots of CG systems. X-axis is distance from 0 to 10 nm perpendicular to nanodisc normal and Y-axis is distance from -5 to 5 nm parallel to nanodisc normal. A plane was fitted to all DMPC beads and LCAT's position relative to it was determined with the same method as described in reference [1]. An illustrative superimposed image is included. The colorbar is normalized by the maximum bin count of apoE helix 1. Systems with less than 10 % peptide dissociation are marked with a dashed red border. The $R^2$ between the dissociation values of non-22A based peptides in non-cancelled systems and the maximum bin counts was 0.38 (positive correlation). In other words, in systems with less peptides on the nanodisc LCAT's movement was more restricted, likely due to entropic forces driving LCAT onto the lipid bilayer. With a peptide rich surface, LCAT is able to sample different angles of the nanodisc perimeter more effectively.

1: Giorgi L, Niemelä A, Kumpula E-P, Natri O, Parkkila P, Huiskonen JT, Koivuniemi A: Mechanistic Insights into the Activation of Lecithin–Cholesterol Acyltransferase in Therapeutic Nanodiscs Composed of Apolipoprotein A-I Mimetic Peptides and Phospholipids. Mol Pharmaceutics 19: 4135-4148, 2022

**Table S1**

| ID | peptide sequence |
|---|---|
| A1 | LKLLDNWDSVTSTFSKLREQLG |
| A2 | PVTQEFWDNLEKETEGLRQEMS |
| A3 | PYLDDFQKKWQEEMELYRQKVE |
| A4 | PLRAELQEGARQKLHELQEKLS |
| A5 | PLGEEMRDRARAHVDALRTHLA |
| A6 | PYSDELRQRLAARLEALKENGG |
| A7 | ARLAEYHAKATEHLSTLSEKAK |
| A8 | PVLESFKVSFLSALEEYTKKLNTQ |
| E1 | ALMDETMKELKAYKSELEEQLT |
| E2 | PVAEETRARLSKELQAAQARLG |
| E3 | ADMEDVCGRLVQYRGEVQAMLG |
| E4 | QSTEELRVRLASHLRKLRKRLL |
| E5 | RDADDLQKRLAVYQAGAREGAE |
| E6 | RGLSAIRERLGPLVEQGRVRAA |
| E7 | TVGSLAGQPLQERAQAWGERLR |
| E8 | ARMEEMGSRTRDRLDEVKEQVA |
| L1 | PVLDLFRELLNELLEALKQKLK |
| L2 | PVLDLFQELLNELLEALKQKLK |
| R1 | PVLDLFDELLNELLEALKQKLK |
| R2 | PVLDLFHELLNELLEALKQKLK |
| R3 | PVLDLFCELLNELLEALKQKLK |
| R4 | PVLDLFLELLNELLEALKQKLK |
| R5 | PVLDLFKELLNELLEALKQKLK |
| M1 | PVLNLFRELLNELLEALKQKLK |
| M2 | PVLNLFQELLNELLEALKQKLK |
| N1 | EMRDRARAHVDALRTHLAPYSD |
| N2-3 | AHVDALRTHLAPYSDELRQRLA |
| N4 | RFWDYLRWVQTLSEQVQEELLS |
| N5 | TLSEQVQEELLSSQVTQELRAL |
| N6 | PLQERAQAWGERLRARMEEMGS |
| N7 | KLEEQAQQIRLQAEAFQARLKS |
| N8 | PLVEDMQRQWAGLVEKVQAAVG |
| N9 | ATYELQREDRALVDTLKFVTQA |
| N10 | KLKETIQKLSNVLQQVKIKDYF |
| N11 | ETNDKIREVTQRLNGEIQALEL |
| N12 | QALELPQKAEALKLFLEETKAT |
| N13 | ETLEDTRDRMYQMDIQQELQRY |
| N14 | YQMDIQQELQRYLSLVGQVYST |
| N15 | IQQELQRYLSLVGQVYSTLVTY |
| N16 | RHFEKNRNNALDFVTKSYNETK |
| N17 | KTTEVLRNLQDLLQFIFQLIED |
| N18 | NLQDLLQFIFQLIEDNIKQLKE |
| N19 | KFNEFIQNELQEASQELQQIHQ |
| N20 | ELQEASQELQQIHQYIMALREE |
| N21 | IISDYHQQFRYKLQDFSDQLSD |
| N22 | TLEDKARELISRIKQSELSAKM |
| N23 | FLKEFPRLKSELEDNIRRLRAL |
| N24 | ELEDNIRRLRALADGVQKVHKG |
| N25 | QQEEYYRKLAGLVEEMHAAYGK |
| N26 | GARDMWRAYSDMREANYIGSDK |
| N27 | TLSEKERQIKKQTALVELVKHK |

**Table S2**

| protein | ID | hydro-philicity | helix penalty (kJ/mol) | resid start | resid end | sequence |
|---|---|---|---|---|---|---|
| apoA1 | A | 0.255 | 2.012 | 21 | 42 | VLKDSGRDYVSQFEGSALGKQL |
| apoA1 | B | 1.132 | 1.692 | 77 | 98 | KETEGLRQEMSKDLEEVKAKVQ |
| apoA1 | C | 0.623 | 1.693 | 99 | 120 | PYLDDFQKKWQEEMELYRQKVE |
| apoA1 | D | 0.859 | 1.96 | 110 | 131 | EEMELYRQKVEPLRAELQEGAR |
| apoA1 | E | 0.664 | 1.978 | 117 | 138 | QKVEPLRAELQEGARQKLHELQ |
| apoA1 | F | 0.677 | 2.155 | 125 | 146 | ELQEGARQKLHELQEKLSPLGE |
| apoA1 | G | 0.564 | 1.47 | 143 | 164 | PLGEEMRDRARAHVDALRTHLA |
| apoA1 | H | 0.555 | 2.042 | 147 | 168 | EMRDRARAHVDALRTHLAPYSD |
| apoA1 | I | 0.186 | 2.056 | 154 | 175 | AHVDALRTHLAPYSDELRQRLA |
| apoA1 | J | 0.623 | 1.532 | 165 | 186 | PYSDELRQRLAARLEALKENGG |
| apoA1 | K | 0.6 | 1.426 | 166 | 187 | YSDELRQRLAARLEALKENGGA |
| apoA1 | L | -0.159 | 2.205 | 209 | 230 | PALEDLRQGLLPVLESFKVSFL |
| apoA1 | M | -0.145 | 2.305 | 210 | 231 | ALEDLRQGLLPVLESFKVSFLS |
| apoE | A | 0.273 | 2.292 | 10 | 31 | PEPELRQQTEWQSGQRWELALG |
| apoE | B | -0.136 | 1.8 | 32 | 53 | RFWDYLRWVQTLSEQVQEELLS |
| apoE | C | 0.155 | 1.553 | 42 | 63 | TLSEQVQEELLSSQVTQELRAL |
| apoE | D | 0.541 | 1.249 | 84 | 105 | PVAEETRARLSKELQAAQARLG |
| apoE | E | 0.641 | 1.297 | 128 | 149 | QSTEELRVRLASHLRKLRKRLL |
| apoE | F | 0.795 | 1.572 | 150 | 171 | RDADDLQKRLAVYQAGAREGAE |
| apoE | G | -0.086 | 2.344 | 183 | 204 | PLVEQGRVRAATVGSLAGQPLQ |
| apoE | H | 0.618 | 1.415 | 202 | 223 | PLQERAQAWGERLRARMEEMGS |
| apoE | I | 1.014 | 1.671 | 209 | 230 | AWGERLRARMEEMGSRTRDRLD |
| apoE | J | 0.482 | 1.177 | 242 | 263 | KLEEQAQQIRLQAEAFQARLKS |
| apoE | K | 0.036 | 1.998 | 252 | 273 | LQAEAFQARLKSWFEPLVEDMQ |
| apoE | L | -0.009 | 1.736 | 267 | 288 | PLVEDMQRQWAGLVEKVQAAVG |
| apoE | M | -0.027 | 1.826 | 268 | 289 | LVEDMQRQWAGLVEKVQAAVGT |
| apoE | N | -0.032 | 2.914 | 278 | 299 | GLVEKVQAAVGTSAAPVPSDNH |
| LCAT | A | -0.236 | 2.759 | 74 | 95 | CWIDNTRVVYNRSSGLVSNAPG |
| LCAT | B | 0.705 | 2.83 | 134 | 155 | VRDETVRAAPYDWRLEPGQQEE |
| LCAT | C | 0.35 | 2.222 | 146 | 167 | WRLEPGQQEEYYRKLAGLVEEM |
| LCAT | D | 0.277 | 1.631 | 152 | 173 | QQEEYYRKLAGLVEEMHAAYGK |
| LCAT | E | 0.073 | 2.747 | 238 | 259 | KLKEEQRITTTSPWMFPSRMAW |
| LCAT | F | -0.327 | 1.978 | 274 | 295 | TGRDFQRFFADLHFEEGWYMWL |
| apoB100 | A | 0.455 | 2.246 | 12 | 33 | CPKDATRFKHLRKYTYNYEAES |
| apoB100 | B | -0.059 | 2.339 | 53 | 74 | VELEVPQLCSFILKTSQCTLKE |
| apoB100 | C | 0.077 | 3.594 | 107 | 128 | AIPEGKQVFLYPEKDEPTYILN |
| apoB100 | D | -0.064 | 2.153 | 142 | 163 | ETEEAKQVLFLDTVYGNCSTHF |
| apoB100 | E | 0.432 | 2.989 | 178 | 199 | TERDLGQCDRFKPIRTGISPLA |
| apoB100 | F | 0.173 | 1.519 | 220 | 241 | YTLDAKRKHVAEAICKEQHLFL |
| apoB100 | G | 0.109 | 2.053 | 442 | 463 | YLMEQIQDDCTGDEDYTYLILR |
| apoB100 | H | 0.791 | 3.07 | 508 | 529 | EPKDKDQEVLLQTFLDDASPGD |
| apoB100 | I | -0.141 | 2.076 | 785 | 806 | MIGEVIRKGSKNDFFLHYIFME |
| apoB100 | J | -0.314 | 2.691 | 863 | 884 | IIPDFARSGVQMNTNFFHESGL |

| | | | | | | |
|---|---|---|---|---|---|---|
| apoB100 | K | -0.373 | 2.606 | 928 | 949 | PLIENRQSWSVCKQVFPGLNYC |
| apoB100 | L | 0.477 | 1.812 | 980 | 1001 | PTGEIEQYSVSATYELQREDRA |
| apoB100 | M | 0.25 | 1.701 | 991 | 1012 | ATYELQREDRALVDTLKFVTQA |
| apoB100 | N | -0.2 | 2.968 | 1284 | 1305 | KMLETVRTPALHFKSVGFHLPS |
| apoB100 | O | -0.164 | 2.016 | 1577 | 1598 | ADYESLRFFSLLSGSLNSHGLE |
| apoB100 | P | 0.45 | 2.07 | 1961 | 1982 | NNNEYSQDLDAYNTKDKIGVEL |
| apoB100 | Q | 0.436 | 2.403 | 2017 | 2038 | DAVEKPQEFTIVAFVKYDKNQD |
| apoB100 | R | 0.114 | 2.542 | 2031 | 2052 | VKYDKNQDVHSINLPFFETLQE |
| apoB100 | S | -0.1 | 1.869 | 2045 | 2066 | PFFETLQEYFERNRQTIIVVLE |
| apoB100 | T | 0.232 | 1.837 | 2052 | 2073 | EYFERNRQTIIVVLENVQRNLK |
| apoB100 | U | 0.023 | 1.895 | 2063 | 2084 | VVLENVQRNLKHINIDQFVRKY |
| apoB100 | V | 0.105 | 1.577 | 2234 | 2255 | QIQEKLQQLKRHIQNIDIQHLA |
| apoB100 | W | 0.25 | 1.685 | 2319 | 2340 | ERYEVDQQIQVLMDKLVELAHQ |
| apoB100 | X | 0.223 | 1.764 | 2342 | 2363 | KLKETIQKLSNVLQQVKIKDYF |
| apoB100 | Y | 0.659 | 2.122 | 2401 | 2422 | KSFDYHQFVDETNDKIREVTQR |
| apoB100 | Z | 0.591 | 1.799 | 2411 | 2432 | ETNDKIREVTQRLNGEIQALEL |
| apoB100 | AA | 0.45 | 2.108 | 2415 | 2436 | KIREVTQRLNGEIQALELPQKA |
| apoB100 | AB | 0.323 | 1.761 | 2428 | 2449 | QALELPQKAEALKLFLEETKAT |
| apoB100 | AC | -0.455 | 1.654 | 2452 | 2473 | VYLESLQDTKITLIINWLQEAL |
| apoB100 | AD | 0.655 | 1.743 | 2487 | 2508 | ETLEDTRDRMYQMDIQQELQRY |
| apoB100 | AE | -0.373 | 1.886 | 2497 | 2518 | YQMDIQQELQRYLSLVGQVYST |
| apoB100 | AF | -0.627 | 1.942 | 2501 | 2522 | IQQELQRYLSLVGQVYSTLVTY |
| apoB100 | AG | 0.577 | 1.98 | 3179 | 3200 | RHFEKNRNNALDFVTKSYNETK |
| apoB100 | AH | -0.136 | 3.84 | 3211 | 3232 | SHDELPRTFQIPGYTVPVVNVE |
| apoB100 | AI | 0.318 | 1.793 | 3345 | 3366 | SVIDALQYKLEGTTRLTRKRGL |
| apoB100 | AJ | 0.218 | 2.359 | 3585 | 3606 | DFPDLGQEVALNANTKNQKIRW |
| apoB100 | AK | 0.164 | 2.327 | 3935 | 3956 | GKYEGLQEWEGKAHLNIKSPAF |
| apoB100 | AL | 0.586 | 1.84 | 4020 | 4041 | ESDEETQIKVNWEEEAASGLLT |
| apoB100 | AM | 0.195 | 1.707 | 4116 | 4137 | EWKDKAQNLYQELLTQEGQASF |
| apoB100 | AN | -0.05 | 1.811 | 4264 | 4285 | KTTEVLRNLQDLLQFIFQLIED |
| apoB100 | AO | -0.009 | 1.649 | 4271 | 4292 | NLQDLLQFIFQLIEDNIKQLKE |
| apoB100 | AP | 0.168 | 1.727 | 4331 | 4352 | KFNEFIQNELQEASQELQQIHQ |
| apoB100 | AQ | 0.236 | 1.412 | 4339 | 4360 | ELQEASQELQQIHQYIMALREE |
| apoB100 | AR | -0.045 | 2.193 | 4343 | 4364 | ASQELQQIHQYIMALREEYFDP |
| apoB100 | AS | 0.109 | 1.967 | 4460 | 4481 | IISDYHQQFRYKLQDFSDQLSD |
| apoB100 | AT | -0.482 | 1.742 | 4486 | 4507 | FIAESKRLIDLSIQNYHTFLIY |
| apoD | A | 0.105 | 2.002 | 34 | 55 | TTFENGRCIQANYSLMENGKIK |
| apoF | A | -0.077 | 2.662 | 7 | 28 | CENEKEQAVHNVVQLLPGVGTF |
| apoF | B | 0.455 | 1.755 | 45 | 66 | KARERGRDGAIDLGYDLLMTMA |
| apoF | C | 0.45 | 1.86 | 109 | 130 | TTKEGLRAISDVSDLEETTTLA |
| apoH | A | 0.355 | 2.678 | 289 | 310 | SYTEDAQCIDGTIEVPKCFKEH |
| apoL1 | A | 0.782 | 1.748 | 72 | 93 | AAAELPRNEADELRKALDNLAR |
| apoL1 | B | 0.877 | 1.321 | 79 | 100 | NEADELRKALDNLARQMIMKDK |
| apoL1 | C | 0.177 | 2.391 | 101 | 122 | NWHDKGQQYRNWFLKEFPRLKS |
| apoL1 | D | 0.645 | 1.905 | 113 | 134 | FLKEFPRLKSELEDNIRRLRAL |
| apoL1 | E | 0.714 | 1.761 | 123 | 144 | ELEDNIRRLRALADGVQKVHKG |

| | | | | | | |
|---|---|---|---|---|---|---|
| apoL1 | F | -0.109 | 2.239 | 133 | 154 | ALADGVQKVHKGTTIANVVSGS |
| apoL1 | G | -0.341 | 2.426 | 298 | 319 | SILEMSRGVKLTDVAPVSFFLV |
| apoM | A | 0.173 | 3.338 | 22 | 43 | QCPEHSQLTTLGVDGKEFPEVH |
| apoM | B | -0.209 | 3.661 | 133 | 154 | MLNETGQGYQRFLLYNRSPHPP |
| apo(A) | A | 0.032 | 2.429 | 1365 | 1386 | GVQDCYRGDGQSYRGTLSTTIT |
| apo(A) | B | -0.005 | 3.051 | 1726 | 1747 | AAQEPHRHSTFIPGTNKWAGLE |
| apo(A) | C | 0.032 | 2.01 | 1870 | 1891 | QEIEVSRLFLEPTQADIALLKL |
| apo(A) | D | -0.291 | 2.57 | 1877 | 1898 | LFLEPTQADIALLKLSRPAVIT |
| apo(A) | E | 0.514 | 3.078 | 1962 | 1983 | RGTDSCQGDSGGPLVCFEKDKY |
| albumin | A | 0.768 | 3.54 | 92 | 113 | AKQEPERNECFLQHKDDNPNLP |
| albumin | B | -0.495 | 2.523 | 138 | 159 | YLYEIARRHPYFYAPELLFFAK |
| albumin | C | 0.118 | 2.059 | 164 | 185 | AFTECCQAADKAACLLPKLDEL |
| albumin | D | 1.041 | 1.463 | 180 | 201 | PKLDELRDEGKASSAKQRLKCA |
| albumin | E | -0.209 | 2.015 | 330 | 351 | FLYEYARRHPDYSVVLLLRLAK |
| albumin | F | 0.227 | 2.384 | 379 | 400 | PLVEEPQNLIKQNCELFEQLGE |
| albumin | G | 0.332 | 2.554 | 422 | 443 | TLVEVSRNLGKVGSKCCKHPEA |
| albumin | H | 0.191 | 2.735 | 439 | 460 | KHPEAKRMPCAEDYLSVVLNQL |
| albumin | I | 0.714 | 1.498 | 515 | 536 | TLSEKERQIKKQTALVELVKHK |
| prothrombin | A | 0.605 | 1.984 | 3 | 24 | TFLEEVRKGNLERECVEETCSY |
| prothrombin | B | 0.359 | 2.144 | 45 | 66 | TACETARTPRDKLAACLEGNCA |
| prothrombin | C | -0.141 | 3.263 | 170 | 191 | CVPDRGQQYQGRLAVTTHGLPC |
| prothrombin | D | 0.532 | 2.008 | 260 | 281 | LDEDSDRAIEGRTATSEYQTFF |
| prothrombin | E | 0.318 | 2.617 | 492 | 513 | VCKDSTRIRITDNMFCAGYKPD |
| prothrombin | F | 0.932 | 3.484 | 511 | 532 | KPDEGKRGDACEGDSGGPFVMK |
| apoC1 | A | 0.745 | 1.329 | 17 | 38 | TLEDKARELISRIKQSELSAKM |
| SAA1 | A | 0.345 | 1.64 | 9 | 30 | EAFDGARDMWRAYSDMREANYI |
| SAA1 | B | 0.495 | 1.684 | 13 | 34 | GARDMWRAYSDMREANYIGSDK |
| SAA1 | C | 0.491 | 1.761 | 60 | 81 | DARENIQRFFGHGAEDSLADQA |
| SAA1 | D | 0.409 | 3.645 | 81 | 102 | AANEWGRSGKDPNHFRPAGLPE |
| SAA2 | A | 0.345 | 1.64 | 9 | 30 | EAFDGARDMWRAYSDMREANYI |
| SAA2 | B | 0.495 | 1.684 | 13 | 34 | GARDMWRAYSDMREANYIGSDK |
| SAA2 | C | 0.65 | 1.64 | 60 | 81 | NARENIQRLTGRGAEDSLADQA |
| SAA4 | A | -0.259 | 1.956 | 6 | 27 | FFKEALQGVGDMGRAYWDIMIS |
| SAA4 | B | -0.045 | 2.128 | 13 | 34 | GVGDMGRAYWDIMISNHQNSNR |
| SAA4 | C | 0.009 | 2.355 | 40 | 61 | GNYDAAQRGPGGVWAAKLISRS |
| SAA4 | D | 1.141 | 3.634 | 89 | 110 | KAEEWGRSGKDPDRFRPDGLPK |

**Figure S1**

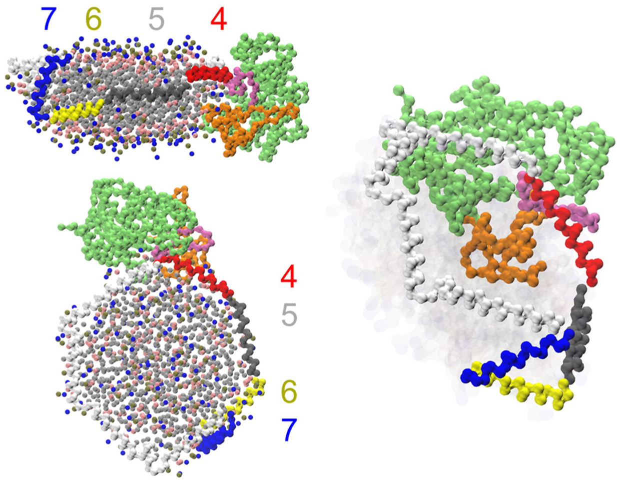


**Figure S2**

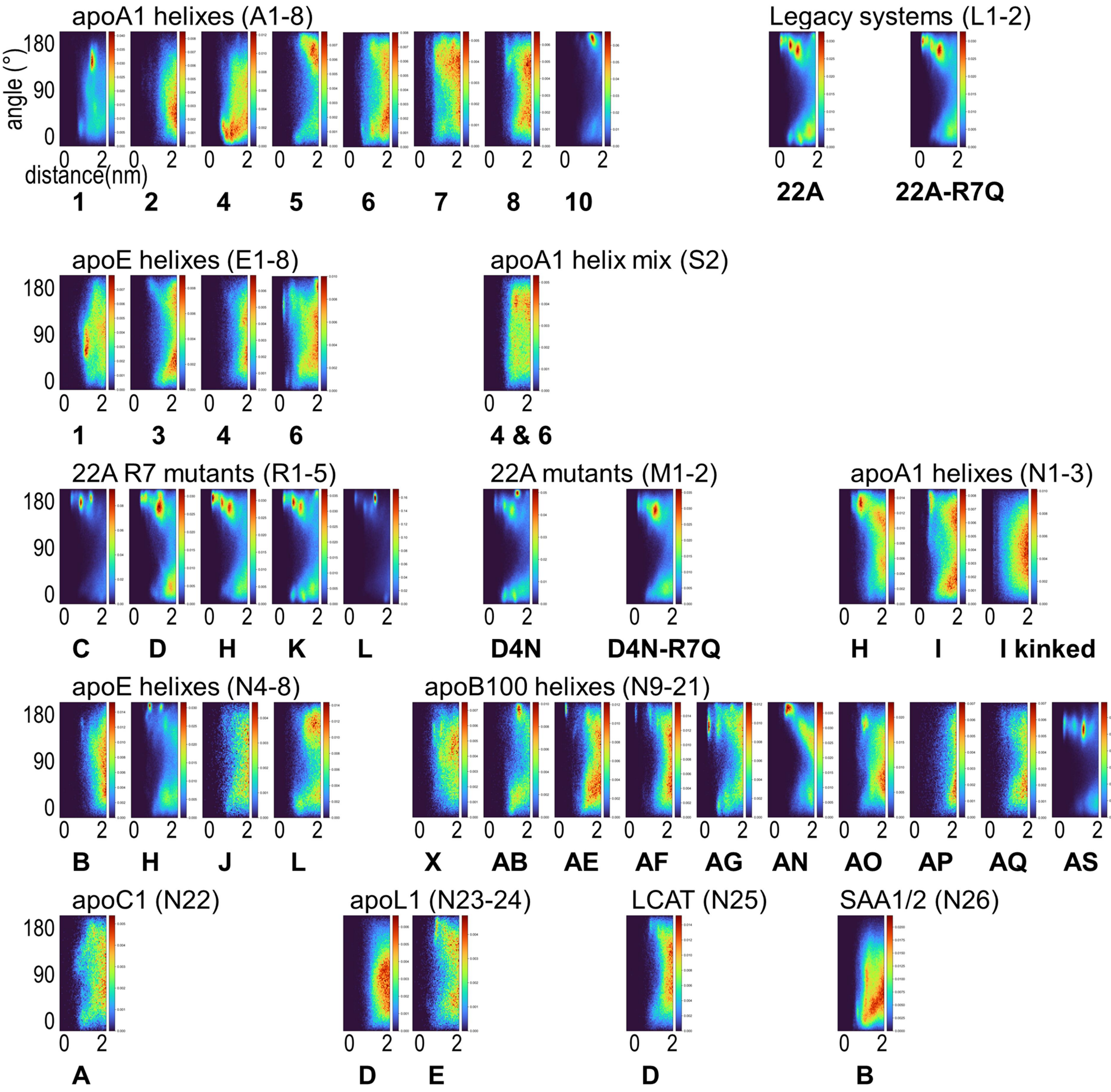

**Figure S3**

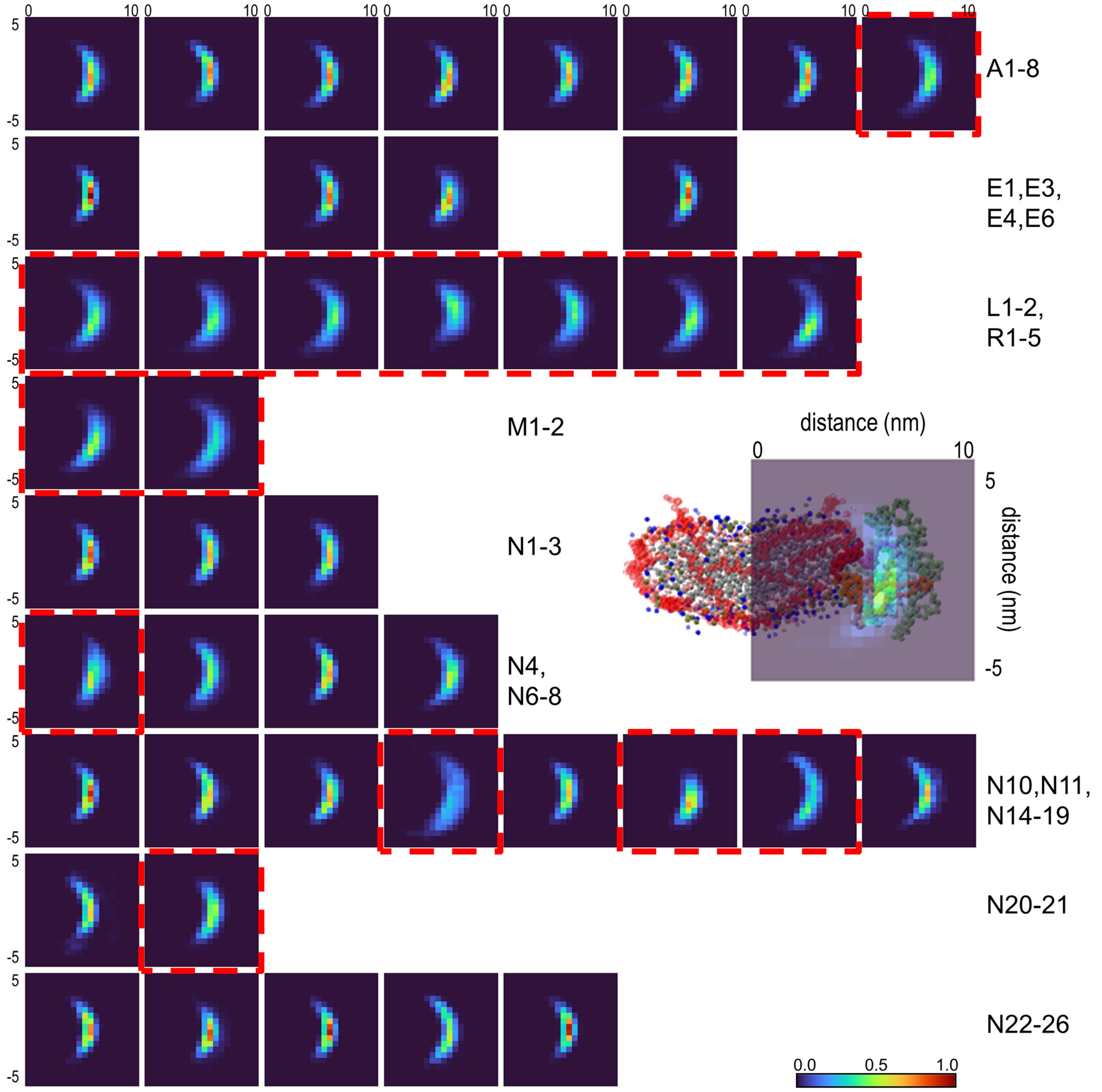
A1-8
E1,E3,
E4,E6
L1-2,
R1-5
M1-2
N1-3
N4,
N6-8
N10,N11,
N14-19
N20-21
N22-26
distance (nm)
distance (nm)
0.0
0.5
1.0
